# Developmental exposure to pesticide contaminated food impedes bumblebee brain growth predisposing adults to become poorer learners

**DOI:** 10.1101/690602

**Authors:** Dylan B. Smith, Andres N. Arce, Ana Ramos Rodrigues, Philipp H. Bischoff, Daisy Burris, Farah Ahmed, Richard J. Gill

## Abstract

Understanding the risk to biodiversity from pesticide exposure is a global priority. For bees, an understudied step in evaluating pesticide risk is understanding how pesticide contaminated foraged food brought back to the colony can affect developing individuals. Provisioning bumblebee colonies with pesticide (neonicotinoid) treated food, we investigated how exposure during two key developmental phases (brood and/or early-adult), impacted brain growth and assessed the consequent effects on adult learning behaviour. Using micro-computed tomography (µCT) scanning and 3D image analysis, we compared brain development for multiple neuropils in workers 3 and 12-days post-emergence. Mushroom body calyces were the neuropils most affected by exposure during either of the developmental phases, with both age cohorts showing smaller structural volumes. Critically, reduced calyces’ growth in pesticide exposed workers was associated with lower responsiveness to a sucrose reward and impaired learning performance. Furthermore, the impact from brood exposure appeared irrecoverable despite no exposure during adulthood.

## Introduction

Insect pollinator declines are of worldwide concern (Hallmann et al., 2017; Potts et al., 2016; Vanbergen, 2013), and safeguarding this important functional group and ecosystem service provider requires a deep understanding of the driving factors (Gill et al., 2016; Goulson et al., 2015). Social bees, such as bumblebees, honeybees and stingless bees are important insect pollinators, and the threat posed by pesticide exposure is a widespread issue (Brittain and Potts, 2011; Desneux et al., 2007; Woodcock et al., 2017). Pesticide residues have been found inside colonies across the globe (Calatayud-Vernich et al., 2018; Daniele et al., 2018; Mitchell et al., 2017; Mullin et al., 2010; Valdovinos-Flores et al., 2017), raising concerns as to how the prevalence of such chemicals in the environment could affect colony development (Gill et al., 2012; Pohorecka et al., 2017; Whitehorn et al., 2012). For instance, controlled exposure experiments and field studies investigating exposure to neonicotinoid pesticides have reported reduced colony growth and sexual output (Arce et al., 2017; Baron et al., 2017; Gill et al., 2012; Leza et al., 2018; Rundlöf et al., 2015; Tsvetkov et al., 2017; Whitehorn et al., 2012). Such colony level effects are likely to be caused by exposure impairing worker behaviour, with cumulative effects across workers leading to a functionally weakened colony (Bryden et al., 2013; Crall et al., 2018). One possibility is that neonicotinoids, being a neurotoxic pesticide, affect neuronal processes important for cognitive and learning abilities (Decourtye et al., 2004; Siviter et al., 2018b) translating to impaired colony tasks (Feltham et al., 2014; Fischer et al., 2014; Gill and Raine, 2014).

With neurotoxic pesticide residues frequently reported in the pollen and nectar brought back by foragers (Botias et al., 2015; Chauzat et al., 2006; David et al., 2016; Kasiotis et al., 2014; Pohorecka et al., 2012), individuals developing and residing in the colony are likely to be chronically exposed to these compounds (Pohorecka et al., 2017). A possibility, therefore, is that tissue development, such as the central nervous system, is impeded. For example, honeybees reared under sub-optimal environmental conditions have exhibited reduced brain volumetric growth and altered neuronal architecture (Groh et al., 2004; Steijven et al., 2017). Impeded brain development and structural plasticity may impact on behaviours such as learning ability, that require detection, assimilation and processing of sensory input from the environment (Cabirol et al., 2018; Chittka, 2017; Galizia et al., 2011). Knowledge of how pesticide contaminated food inside bee colonies can affect individual physiological development however, is limited (Gregorc et al., 2012; Wu et al., 2012, 2011). Moreover, there has been an urgent call for research linking how potential pesticide induced impairment to brain development can translate to task performance as later adults (Siviter et al., 2018b; Tan et al., 2015; Tomé et al., 2012; Yang et al., 2012). We directly address this call by investigating how developing bumblebees exposed to a neonicotinoid pesticide via treated provisioned food may alter brain development and link this to effects on associative learning behaviour.

To experimentally test the effect of pesticide exposure on individual development we needed to first consider the level of developmental plasticity in behaviour and brain growth. In social bees, worker maturation can correlate with stereotyped behavioural changes (Goulson, 2010; Johnson, 2010), and increased brain neuropil volumes (functional structures) (Durst et al., 1994; Galizia et al., 2011; Li et al., 2017; Winnington et al., 1996; Withers et al., 1993). However, it has been recognised that dissecting innate effects of age (experience independent change) from co-varying cumulative increases in sensory input (experience dependent change) on behaviour and brain development is difficult (Fahrbach et al., 1998; Jones et al., 2013; Maleszka et al., 2009; Riveros and Gronenberg, 2010). To distinguish the effects of pesticide exposure from variation caused by other interacting factors, we therefore: i) attempted to standardise experience and sensory input across tested workers, ii) tested workers of controlled age, and iii) compared between young and old age cohorts. Furthermore, by studying two main developmental stages, such as brood (larval & pupal) and early adulthood, here we reveal which development phase is more vulnerable to pesticide exposure, and whether developmental plasticity in bee brains (Farris et al., 2001; Galizia et al., 2011; Riveros and Gronenberg, 2010; Withers et al., 1995) allows recovery during an unexposed later stage.

Despite the functional importance of non-*Apis* bees (Brittain et al., 2013; Garibaldi et al., 2013; Gill et al., 2016) and possible differences in pesticide sensitivity relative to honeybees (Cresswell et al., 2012; Heard et al., 2017; Piiroinen and Goulson, 2016; Rundlöf et al., 2015; Woodcock et al., 2016), empirical tests on non-*Apis* bees looking at the physiological response to stress, such as postembryonic neuronal development, are limited. Here we studied the response of the bumblebee, *Bombus terrestris,* a species: i) that can be reared under controlled laboratory conditions; ii) for which learning performance assays have been developed (Muth and Leonard, 2019; Riveros and Gronenberg, 2009; Siviter et al., 2018b; Smith and Raine, 2014); iii) individuals can be exposed to pesticides within the social colony environment rather than in isolation (Gill et al., 2012; Maleszka et al., 2009; Whitehorn et al., 2012). Here, we chronically exposed cohorts of workers, reared inside their natal colonies, to a 5ppb concentration of the neonicotinoid imidacloprid via provisioned sugar solution (40% sucrose) during two different developmental stages: a) before and b) after adult eclosion from the pupal case. We investigated how the link between brain growth and learning behaviour may be affected in workers exposed during: brood development (*pre-eclosion*), early adulthood (*post-eclosion*) and both these developmental periods (*continual*), comparing each to unexposed (*control*) workers. For each treatment we tested adult workers at 3 or 12-days after eclosion (Figure 1).

**Figure 1.**
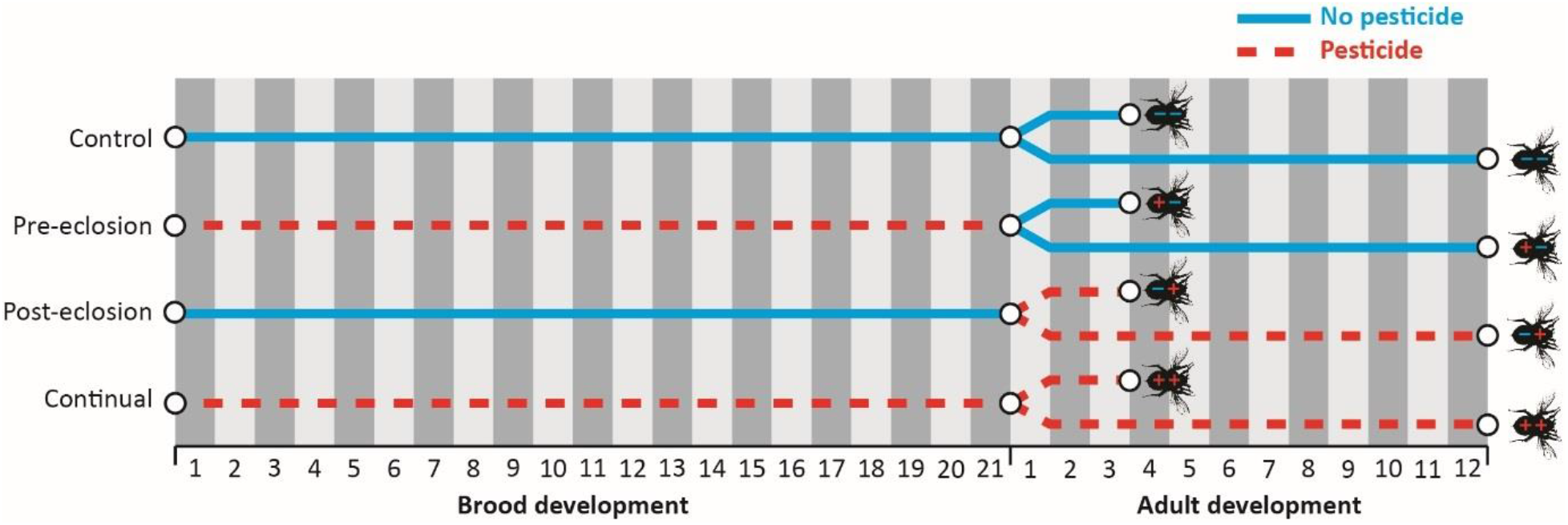
Graphic showing exposure periods for the four colony treatments (*control, pre-eclosion, post eclosion & continual*) and the eight cohorts of individual workers to be tested. Blue solid line represents untreated food (sucrose solution) and red dashed line represents pesticide-treated food. ‘Brood development’ represents the larval and pupal (brood) stages of workers, with ‘Adult development’ representing the number of days after eclosion from the pupal case. Individual bee symbols depict removal of controlled aged adult workers at 3 or 12-days after eclosion for behavioural testing followed by decapitation for µCT scanning of the brain.

Firstly, we tested worker response to sucrose and then on olfactory associative learning performance using the established proboscis extension reflex (PER) conditioning paradigm (Figure S1) (Bitterman et al., 1983; Giurfa and Sandoz, 2012; Laloi et al., 1999; Riveros and Gronenberg, 2009; Sommerlandt et al., 2014), which has previously been used to test pesticide effects on adult learning in honey bees exposed during the larval stage (Tan et al., 2015; Yang et al., 2012), and bumblebees exposed as adults (Piiroinen and Goulson, 2016; Stanley et al., 2015; Tison et al., 2017). Secondly, we employed new advances in micro-computed tomography (µCT) scanning and 3D image analysis to explore the brain *in situ* (within headcase) and enable non-destructive volumetric measurements to a standardised voxel size of 4µm (Figure 2) (Baird and Taylor, 2017; Gutiérrez et al., 2018; Ribi et al., 2008; Smith et al., 2016). We segmented five key neuropils: mushroom bodies (associated with learning), antennal lobes (olfaction), optic lobes - medullas and lobulas (vision), and central body (motor function; Table S1). For the mushroom bodies we segmented the two major components, lobes and calyces, to investigate responses of the different functionally multisensory input and output regions, respectively (Fahrbach, 2006; Heisenberg, 2003). Using a sample size exceeding any other study investigating bee brain morphology to date, we present the first comparative study to directly link how chronic pesticide exposure impacts learning performance by affecting bee brain development.

**Figure 2.**
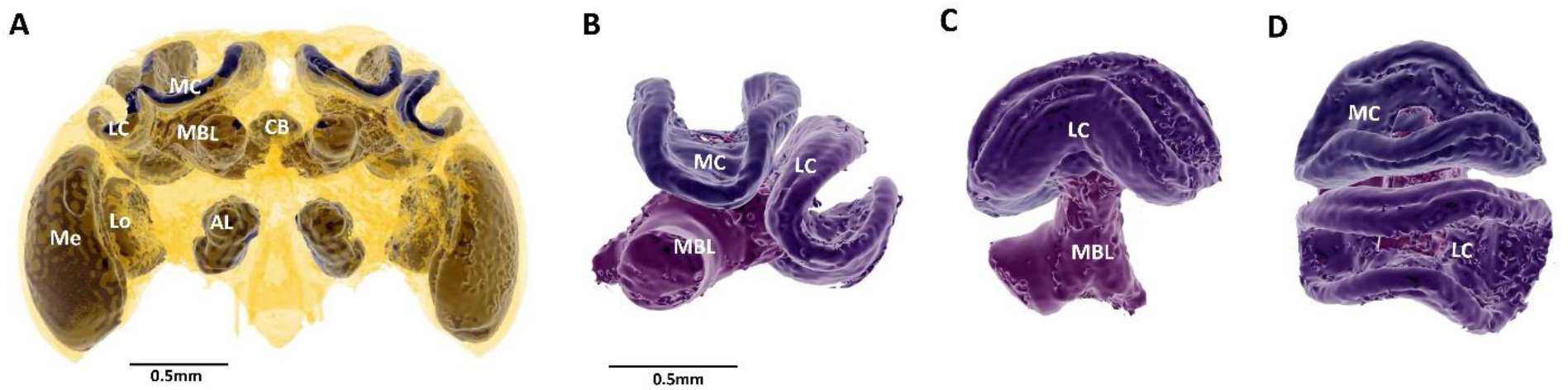
3D rendering of one of the studied bumblebee brains using the µCT imaging method. **a**, Focal neuropils considered in this study shown in dark purple (optic lobes: medulla (Me), lobula (Lo); antennal lobes (AL); central body (CB); mushroom body: calyces (including lateral calyx (LC) & medial calyx (MC) and lobes (MBL)), surrounded by remaining brain tissue in transparent yellow. **b-d,** Isolated 3D structure of the mushroom body which has been rotated to show **b**, frontal, **c**, lateral, and **d**, dorsal views.

## Results

### Responsiveness

Prior to the learning assay, we confirmed whether harnessed workers (n=413; Table S2) showed a PER in response to their antenna being touched by a 50% sucrose solution droplet (Figure S1). We found a significantly higher proportion of 12-day compared to 3-day workers responded (GLM: *age*: z=-4.10, p<0.001) which was consistent across treatments as evidenced by no *age*treatment* effect and the interaction term not being retained in the model (Table S3). We also found consistent negative model estimates for all three pesticide treatments relative to *control*, and detected a significantly lower proportion of responsive workers from *post-eclosion* and *continual* exposed colonies (z=-2.53, p=0.011 & z=-2.40, p=0.016; Figure 3).

**Figure 3.**
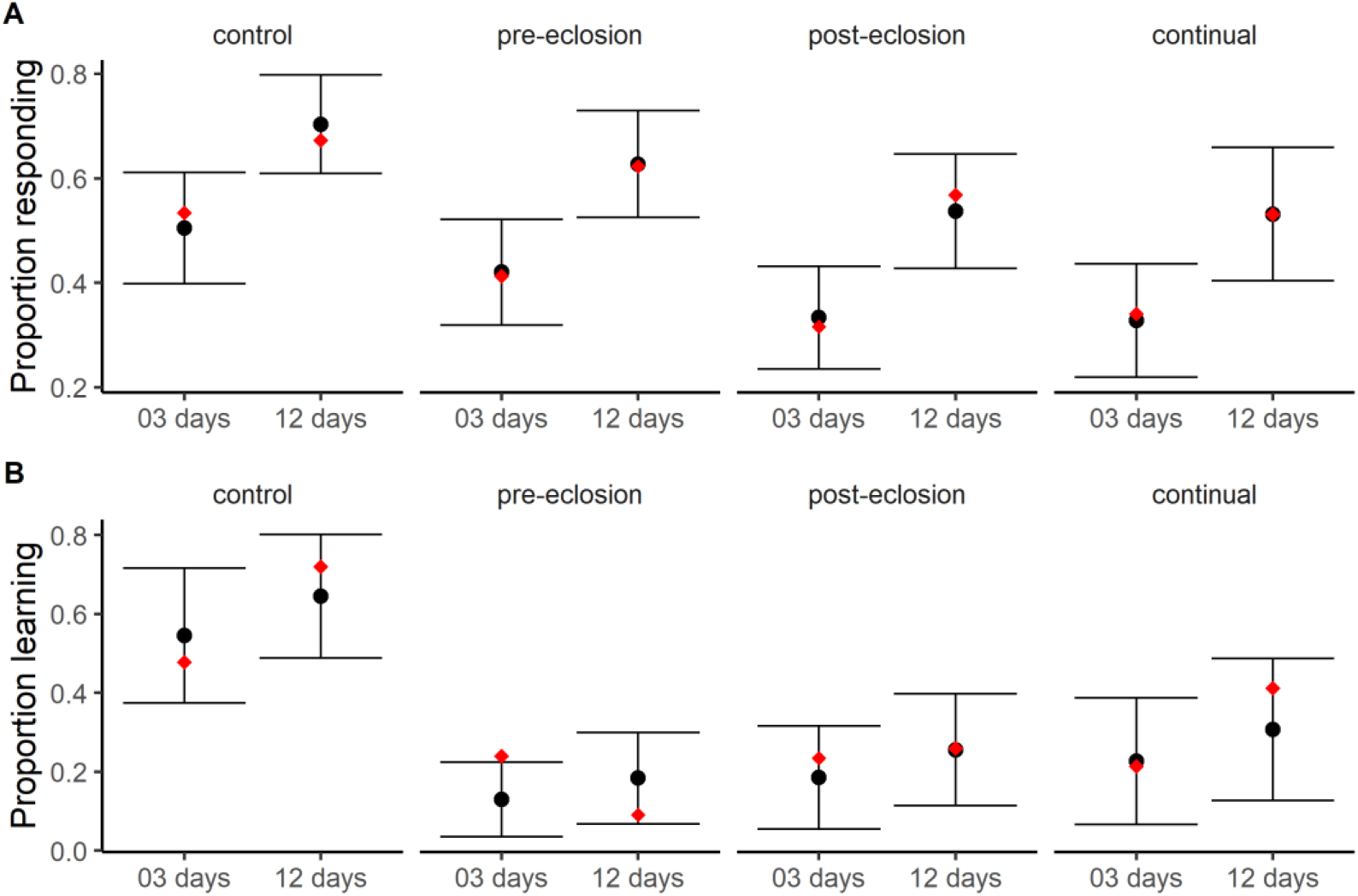
Proportion of responsive workers and proportion demonstrating an olfactory associative learning response using proboscis extension reflex (PER) conditioning between treatments. **A,** Workers exhibiting a PER response when touching the antennae with a sucrose solution droplet prior to the PER conditioning trials; **B,** Learners (workers exhibiting at least one learnt response during the PER conditioning trials). Intersecting circular point represents the estimated model mean taken from back-transformation of the model (binomial GLM) with bars depicting the associated ±95% confidence limits. Red diamond corresponds to the mean value taken from the raw response data.

### Learning

For the responsive workers (n=181; Table S2), we tested each worker’s ability to learn to associate an odour with a sucrose reward by demonstrating a PER response over ten consecutive trials (Figure S1; see methods). Firstly, we categorised workers as either those exhibiting at least one response as ‘learners’ and those showing no learnt response as non-learners. Whilst our model showed a positive estimate for the effect of *age*, unlike responsiveness we did not detect a significant increase which was consistent across treatments as evidenced by no *age*treatment* effect and the interaction term not being retained in the model (Table S3). However, we again detected a strong effect of pesticide exposure relative to the *control*, with each treatment showing a significantly lower proportion of learners (GLM: *pre-eclosion*: z=-4.38, p<0.001; *post-eclosion*: z=-3.49, p<0.001; *continual*: z=-2.78, p<0.01; Figure 3b).

For all individuals classed as learners, we then looked at how the proportion of learnt responses changed over the successive trials (analysis considered trials 2-10, as by definition a naïve worker cannot learn on the first trial). However, because of the strong negative effect of pesticide exposure on passing the responsiveness stage and proportion of learners, sample sizes for each pesticide treatment were significantly reduced. Therefore, given the similarity in responses across pesticide treatments, we pooled all workers from these three treatments and compared them to *control* whilst not distinguishing between 3-and 12-day workers. From this analysis we found that the proportion of responses increased over the trials (GLM polynomial: *p^1^*: t=14.26, p<0.001). This relationship, however, was non-linear and the incremental proportion decreased in rate over the consecutive trials (*p^2^*: t=-2.48, p=0.014; Table S3). This was primarily driven by the significant negative effect of pesticide exposure (t=-2.04, p=0.046), with workers from exposed colonies showing a distinctly lower proportion of learnt responses in the latter few trials relative to *control* (Figure 4).

**Figure 4.**
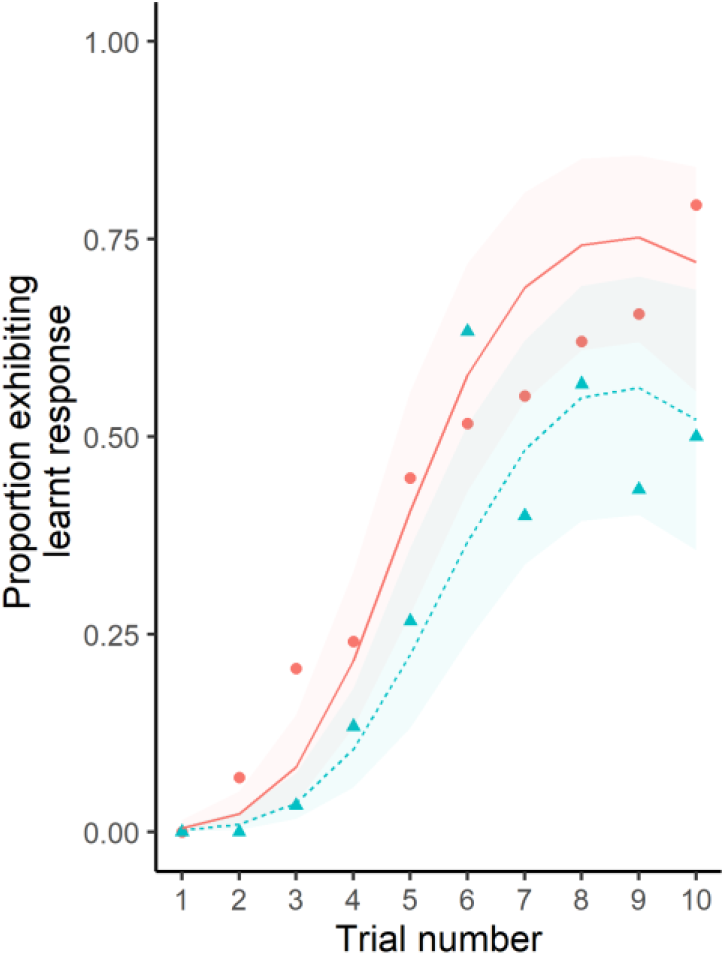
Proportion of workers by trial exhibiting an olfactory conditioned learnt response. Workers from all three pesticide treatments were pooled (blue triangles; n=30 workers) and compared against *control* workers (red circles; n=29), with both age cohorts aggregated per treatment. Lines (blue dashed = pesticide treatment; red solid = control) represent the binomial model (LMER polynomial) estimates over the consecutive trials.

### Brain neuropil volumes

Focusing first on the mushroom body calyces, relative volumes were significantly smaller in workers from all three pesticide exposure treatments compared to *control* (*pre-eclosion*: t=-2.41, p=0.049; *post-eclosion*: t=-3.83, p<0.01; *continual*: t=-2.90, p=0.021; Table S4-5). This was consistent for both 3 and 12-day workers as evidenced by no effect of *age*treatment* and the interaction term not being retained in the model (Table S5). Focusing second on the relative volume of the mushroom body lobes, we again found negative model estimates for all three pesticide treatments relative to the *control*, however unlike the calyces none of these comparisons were detected as significantly lower (Table S5). Analysis of the four other segmented neuropils (central body, antennal lobes, lobulas and medullas) further showed that workers from pesticide treated colonies showed no significant volumetric differences relative to *control*, although we did find consistent negative model estimates for the antennal lobes across all pesticide treatments (Table S6).

### Relationship between mushroom body calyces volume and learning score

For each responsive worker that started the PER conditioned learning assay and for which we had the volume of their mushroom body calyces, we took the total number of demonstrated learnt responses (‘*learning score*’) and investigated the association with relative calyces’ volume as the predictor variable (given this was the neuropil most affected from exposure). As we did for the learning-by-trial data, we pooled all workers from the three pesticide exposure treatments and compared their scores to that of *control* workers when no distinguishing age. From this, we found a significant positive association between relative volume of the calyces and learning score (t=4.51, p<0.001; Figure 6; Figure S2), but this relationship was driven by *control* workers in which larger calyces equated to higher learning score. Pesticide exposed workers, in contrast, showed no clear relationship as supported by the significant negative *volume*treatment* interaction (t=-3.96, p<0.001; Table S7-8). This finding reveals that impaired functioning of the mushroom body in workers from pesticide exposed colonies is not only from reduced volumetric growth, but presumably also from affected physiological composition of the tissue.

**Figure 5.**
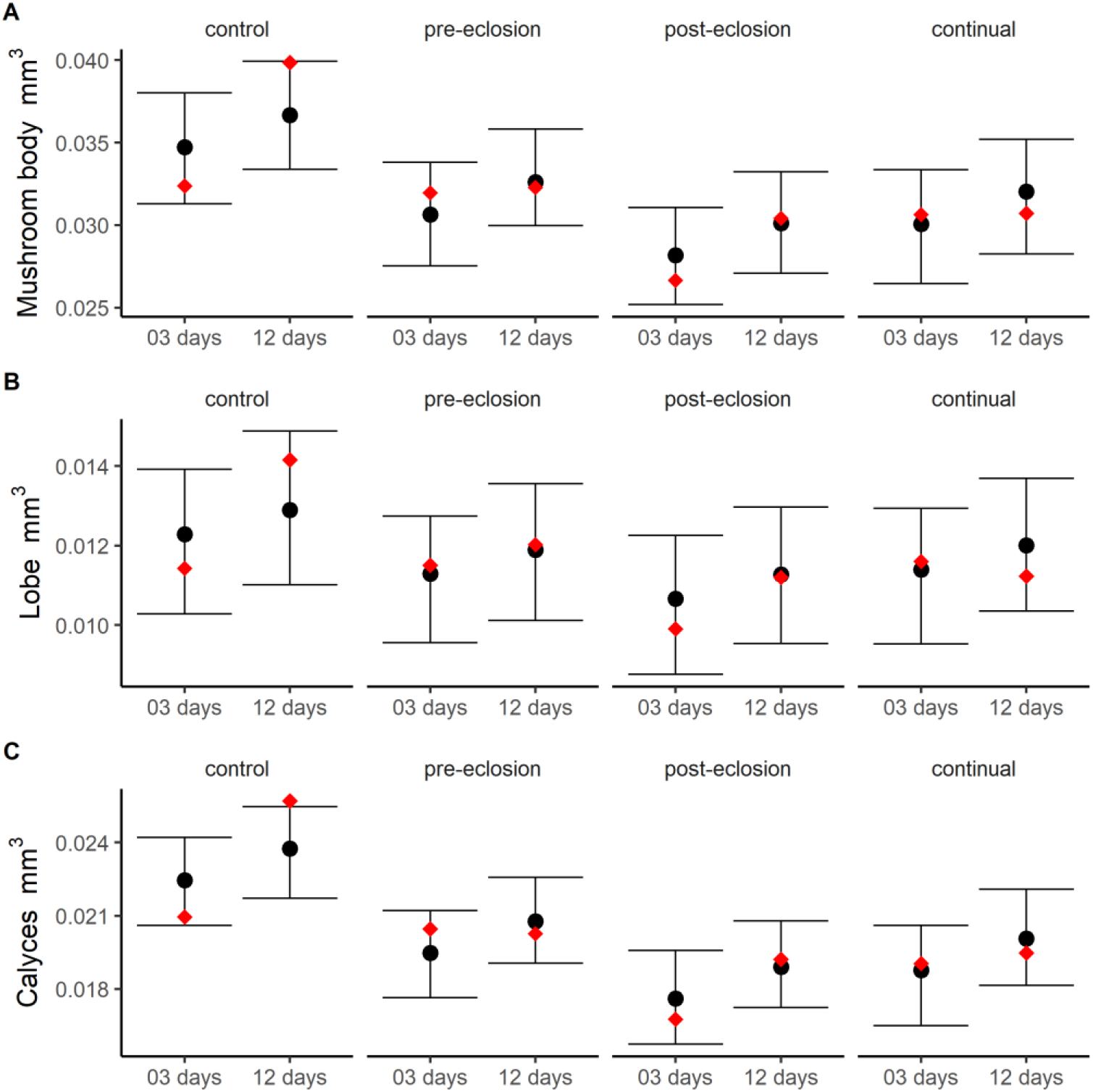
Volumes for A) whole mushroom body, B) mushroom body calyces and C) mushroom body lobes of bumblebee workers. These represent volumes relative to the body size of the worker (absolute volume divided by body size). Intersecting circular point represents the estimated model mean taken from back-transformation of the model (LMER) with bars depicting the associated ±95% confidence limits. Red diamond corresponds to the mean value taken from the raw response data.

**Figure 6.**
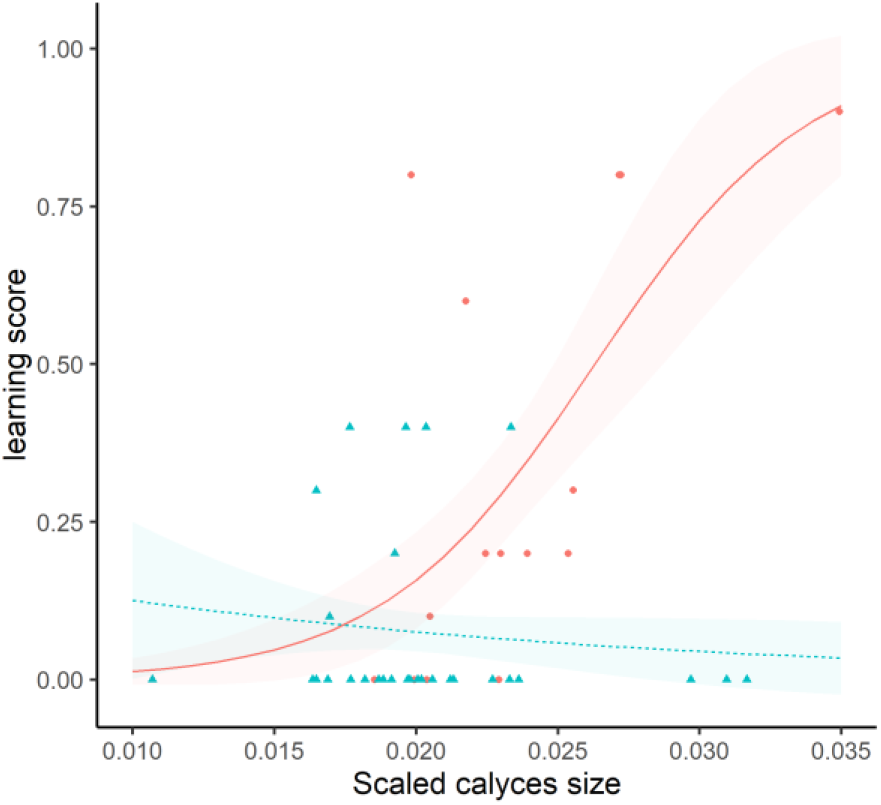
Relative volume of mushroom body calyces plotted against learning score. Calyces volume predicts learning score for *control* workers but not pesticide exposed workers. Workers from all three pesticide treatments were pooled (blue triangles) and compared against *control* workers (red circles), with fitted lines (blue dashed = pesticide treatment; red solid = control) representing binomial model (GLM) estimates.

## Discussion

Our study reveals that worker bumblebees exposed to a neurotoxic pesticide, a neonicotinoid, can affect the developmental plasticity of the brain with reduced volumetric growth manifested not only from exposure as an adult but also during brood development. The effects on adult behaviour and brain physiology from brood exposure (*pre-eclosion*) appeared irrecoverable despite no experimental provision of pesticide treated food during the 12-days of adulthood. Critically, impeded growth of the mushroom body calyces of worker brains from pesticide exposure, was associated with functional impairment as evidenced by reduced responsiveness and poorer olfactory learning behaviour.

### Pesticide exposure during early development affected responsiveness and learning

Neonicotinoid exposure as an adult (*post-eclosion* & *continual*) reduced the proportion of workers responding to a sucrose droplet prior to the PER assay, and reduced olfactory learning performance during the PER assay. These findings contribute to previous studies reporting adult neonicotinoid exposure negatively affecting aspects of responsiveness in honeybees (Aliouane et al., 2009; Démares et al., 2018, 2016) and learning in bumblebees (Stanley et al., 2015). However, a key novelty of our study is that we could compare responses from chronic exposure between age cohorts. Firstly, this revealed that young (3-day) and older (12-day) workers from *post-eclosion* and *continual* exposure colonies were similarly affected despite differences in the number of days of adult exposure. Additionally, despite *pre-eclosion* 3-day adults being exposed for up to 3-weeks during brood development, compared to only three days of exposure for 3-day adults from *post-eclosion* colonies, the degree of impaired learning was again similar.

Together these findings highlight the first 72 hours of adulthood to be critical in behavioural development, and reveals a susceptible developmental window to environmental stress (in this case pesticide exposure) (Sandrock et al., 2014; Wu et al., 2011), reiterating the importance of considering different life-stages when assessing pesticide risks.

Secondly, workers exposed during brood development (*pre-eclosion*), that received no or substantially lower exposure as an adult, exhibited impaired learning performance at a similar reduced level as adult only exposed workers (*post-eclosion*). This indicates a lag-effect from brood exposure on adult learning, highlighting the importance of considering delayed effects of pesticide exposure; a view reinforced by other studies on honeybees (*Apis cerana & A. mellifera*) and a stingless bee (*Melipona quadrifasciata anthidioides*) reporting larvae reared under topical or oral neonicotinoid exposure exhibited negative effects on adult learning and motor function (Tan et al., 2015; Tomé et al., 2012; Yang et al., 2012). More importantly, with 3- and 12-day adults from *pre-eclosion* colonies exhibiting a similar level of impaired learning performance, this reveals that the effects from brood exposure appear irrecoverable even as an unexposed adult.

### Impaired learning in pesticide exposed workers was associated with reduced volumetric growth of the mushroom body calyces

Focusing on the mushroom body calyces, 12-day adult workers from all pesticide exposure treatments possessed smaller mushroom body calyces relative to *control* colonies, and we even found differences in 3-day adult workers from *post-eclosion* and *continual* exposed colonies. With rates of mushroom body development in bumblebees considered to be at its highest during the first 72 hours of adulthood (Jones et al., 2013; Riveros and Gronenberg, 2010), this may explain why in our experiment an effect on calyces volume was detected in just 3-days of adulthood. Furthermore, our finding that 3-day adults from both *pre-eclosion* and *post-eclosion* exposed colonies showed similar reductions in mushroom body volumes, reiterates the apparent vulnerability of brain development during the first 72 hours.

Average volume reductions of the mushroom body calyces and lobes showed a strikingly mirrored pattern to the reduced proportion of learners in each respective pesticide treatment. More tellingly, when relative volume of the calyces was plotted against each respective worker’s learning score, bigger relative brain size equated to better learning performance in *control* workers, but this relationship was not found for pesticde exposed workers. Despite some *control* and pesticide exposure workers possessing similar relative mushroom body volumes, pesticide exposure workers demonstrated a lower learning score indicating impaired neuronal functioning of this brain region. For workers from *post-eclosion* and *continual* exposed colonies this effect could be explained by sublethal neonicotinoid concentration affecting neuronal signalling given its role as a nAChE receptor agonist (Palmer et al., 2013). However, with *pre-eclosion* workers also being affected, this indicates that neonicotinoid exposure might be affecting synaptic development (i.e. proliferation & dendritic outgrowth) in the calyces. Indeed, reduced microglomeruli density has been shown to occur in neonicotinoid exposed honeybees (Peng and Yang, 2016), and density of these structures is correlated with increased learning and memory in bees (Hourcade et al., 2010; Li et al., 2017). Furthermore, the reduced learning performance in *pre-eclosion* workers in our study could stem from impeded neurogenesis where neuronal precursor cells are in some way prevented from giving rise to Kenyon cells in the mushroom bodies, which in honeybees occurs during development before eclosion from the pupal case (Fahrbach et al., 1995; Farris et al., 1999). Alternatively, the size of Kenyon cells could be affected, as has been shown from exposure experiments on bumblebee cell cultures (Wilson et al., 2013).

### Mushroom body calyces were disproportionately affected over the other neuropils

The effect of neonicotinoid exposure was primarily localized to the mushroom body, and even within this structure was manifested more heavily in the calyces than the lobes. Localised variation in plasticity has been shown in bumblebees where foraging experience increased medial but not lateral calyx volume (Riveros and Gronenberg, 2010). The calyces act as multisensory processors fed by afferent neurons, whereas the lobes predominantly function as output regions with efferent neurons, which could explain why calyx volumetric variation is more tightly associated with our measure of learning performance. Antennal lobes are involved in detecting and processing olfactory information (Galizia et al., 1999; Hansson and Anton, 2002; Sachse et al., 1999), developmentally plastic during early adulthood (Jones et al., 2013; Riveros and Gronenberg, 2010), and exhibit reduced neuronal function under nicotinic agonists (Andrione et al., 2016; Barbara et al., 2008; Thany and Gauthier, 2005). Considering pesticide exposed workers performed worse in the olfactory conditioning, we might therefore expect a pattern of impeded growth in the antennal lobes similar to that found for the mushroom body calyces. Whilst in support of this view we found consistent negative model estimates for all three pesticide treatments, unlike the calyces our analysis did not detect a significant effect. Furthermore, we found no consistent reduction in volume of the optic lobes (medullas & lobulas) or central body for pesticide exposed workers.

Possible explanations for the disproportionate effect of neonicotinoid exposure on the mushroom bodies may include: i) nACh receptors targeted by neonicotinoids are found in the highest density in the Kenyon cells of the mushroom bodies (Galizia et al., 2011) and so could affect Kenyon cell proliferation leading to volumetric reductions; ii) The mushroom bodies, in particular the calyces, of social insects have consistently been reported to be highly plastic structures due to their role in learning and memory development as early adults (Cabirol et al., 2018; Farris et al., 2001; Heisenberg, 2003; Jones et al., 2013; Kühn-Bühlmann and Wehner, 2006; Riveros and Gronenberg, 2010). The large amount of neuronal development and re-organisation therefore increases the risk of neurotoxic exposure interfering with this process; iii) Our experimental setup was stimulus deprived and not void, therefore whilst mushroom body volumetric increase is likely to be more experience independent than dependent, we could not rule out investment in olfactory processing to compensate for a lack of visual information (Fahrbach et al., 1998; Jones et al., 2013); iv) the change in growth of non-mushroom body neuropils was simply too subtle for our μCT technology and/or sample sizes to detect.

### Improved behavioural performance and mushroom body growth with age independent of experience

Our study reared workers under a stimulus deprived environment, therefore a positive effect of age is indicative of experience independent age-enhanced learning. For both our measures of responsiveness and learning we found positive estimates for age, with 12-day workers performing better on average than 3-day. This finding contrasts with previous bumblebee studies reporting no effect of age on aspects of learning ability (Riveros and Gronenberg, 2009; Smith and Raine, 2014), but these were carried out in foraging arenas whereby prior experiences could not be fully controlled. Indeed, to our knowledge, there is a lack of studies looking to identify innate age-related growth on bumblebee behaviour as well as on brain growth. Only one histological study on the bumblebee *Bombus impatiens* by Jones and colleagues has shown age-dependent volumetric growth in brain neuropils separate form environmental stimuli (Jones et al., 2013). Their findings suggested c.10% increase in the mushroom body calyces and lobes which interestingly is around half the increase we found in our *control* workers (just over 20%), a difference that perhaps stems from variation in methodological approaches, sample sizes (lower in Jones et al.) or taxonomic variation. However, together this evidence supports an innate increase in neuropil volume over the first 12-days of adulthood, which presumably is important to prepare workers for the complex colony tasks required at this age (Maleszka et al., 2009).

### Implications of our findings for social insects

Our findings that early exposure effects later adult behaviour provides a mechanistic explanation for why reduced colony growth is often detected 2-3 weeks after onset of neonicotinoid exposure (Arce et al., 2017; Gill et al., 2012; Rundlöf et al., 2015; Siviter et al., 2018a; Tsvetkov et al., 2017; Whitehorn et al., 2012), and why reduced colony productivity has been correlated with neonicotinoid treated neighbouring fields (Rundlöf et al., 2015; Woodcock et al., 2016). With eusocial bee colonies having overlapping generations, colonies are reliant on newly emerging cohorts of workers to be effective task performers. If future generations of workers are predisposed to be inefficient functioning cohorts, this could lead to a density dependent build-up of colony level impairment increasing the risk of colony collapse (Bryden et al., 2013). Our results suggest that even if workers were to delay undertaking a task, such as foraging, in attempt to developmentally recover, this strategy may be futile given we saw little adult recovery from 3 to 12 days of adulthood from *pre-eclosion* colonies. Importantly, these effects are unlikely to be exclusively applicable to neonicotinoids as a multitude of other neurotoxic pesticides including the possible neonicotinoid replacements, sulfoxamines and butenolides (Siviter et al., 2018a; Tosi and Nieh, 2019), are likely to end-up inside bee colonies with the potential to influence tissue development in reared bees.

## Materials & methods

### Animal Husbandry

Twenty-two *Bombus terrestris audax* colonies were delivered by a commercial supplier (Agralan Ltd), with colonies possessing a queen and mean (±s.e.m.) of 14.5 ± 1.1 workers on arrival (Table S9) and housed in an aerated plastic box (29 x 22.5 x 13 cm). All colonies were moved to a controlled environment (23°C; 60% humidity) red light room where they remained for the duration of the experiment. Throughout the experiment, colonies were provisioned untreated honeybee collected pollen (supplied by a commercial supplier Agralan Ltd) ad-libitum in a petri dish, and 40/60% sucrose/water solution in a gravity feeder. Food was replenished every two days, and feeders thoroughly cleaned prior to refill (Table S10 for colony consumption). During Phase I (days 1-21; Figure 1; Figure S3), we conducted daily checks of all newly eclosed bees and marked each using a white paint pen (uni Posca, PC.5M 1.8-2.5mm), allowing us to distinguish between newly eclosed workers during Phase II (day 22 onwards) from eclosed workers before this. Colonies were checked daily for males, gynes or dead individuals which were removed and frozen at −20°C.

### Experimental setup

On arrival, colonies were randomly assigned to the four treatments, with no significant difference in the number of workers between treatments (ANOVA: F_3,22_=1.04, p=0.40). Mean worker thorax width was similar between treatments (LMM, p>0.07) with *control* being 4.23mm (range=3.29-5.17), *pre-eclosion* 4.16mm (3.09-5.12), *post-eclosion* 4.28mm (3.14-5.63) and *continual* 4.33mm (3.36-5.34). Monitoring overall development of workers in colonies, we implemented a fully factorial design with our colony treatments comprising a combination of two exposure phases: Phase I encompassing the majority of brood (larval & pupal) development period and Phase II comprising the early adult development period (up to 12 days). Phase I exposure period started two days after colonies arrived and lasted for 21 days approximating development time from an egg or very small larva to adult eclosion (Alford, 1975; Cnaani et al., 2002; Duchateau and Velthuis, 1988). This ensured that all sampled adults will have been exposed/unexposed in a standardised manner during the vast majority of brood development (Figure S3). On the 22^nd^ day Phase II started, during which we checked daily for callow workers (adults recently eclosed from their pupal case) and tagged each with a unique numbered Opalith tag using superglue. On tagging, we randomly assigned half of the workers per colony per day to the 3-day cohort and remaining half to the 12-day cohort, with tag ID used to correctly remove for testing 3 or 12 days later. Tagging period lasted 11 days to provide us with a high number of workers to test (Table S2). This window of opportunity approximates the minimum time of pupal development, in which pupae evacuate their gut and stop feeding (Cnaani et al., 2002), allowing us to standardised pesticide exposure as best possible across all tested workers. Adult workers aged 3 and 12-days after eclosion were chosen to be tested as brain development has been reported to occur both during brood and early adult stages (Farris et al., 2001; Jones et al., 2013).

We applied four treatments to colonies: *control* = phases I & II unexposed (n=5 colonies); *pre-eclosion* = phase I exposed to Imidacloprid, phase II unexposed (n = 6); *post-eclosion* = phase I unexposed, phase II exposed to Imidacloprid (n = 6); *continual* = phases I & II exposed to Imidacloprid (n = 5). The neonicotinoid imidacloprid was used as: i) it is widely used across the globe (Casida, 2018; Cressey, 2017; Mitchell et al., 2017; Zhang, 2018); ii) it targets nAChE receptors found in insect brains (Jeschke and Nauen, 2008; Palmer et al., 2013); iii) exposure has been shown to affect bee foraging and navigation known to be reliant on learning ability and working memory (Feltham et al., 2014; Fischer et al., 2014; Gill and Raine, 2014; Samuelson et al., 2016; Stanley and Raine, 2016). The imidacloprid treated sucrose solution provisioned to the colony was made from a primary stock solution (1mg/ml) consisting of 100mg of imidacloprid (powder; grade: PESTANAL®, analytical standard; brand: Fluka) dissolved in 100ml of acetone. An aliquot was then added to a 40/60% sucrose/water solution to produce a 5ppb imidacloprid solution of required volume. A *control* sucrose solution was made by repeating this process but a same aliquot volume of pure acetone.

### Assessing olfactory learning performance using proboscis extension reflex (PER) conditioning

The proboscis extension reflex (PER) conditioning paradigm we implemented was adapted from a previously reported setup on bumblebees (Stanley et al., 2015). On removal from the colony, workers were harnessed (between 13:00-14:00) using a modified 2ml centrifuge tube and a split pin yoke, under natural light in the lab and left for 2hrs to settle (Figure S1 for harness setup). All bees were then fed to satiety using a Gilmont® syringe to present 40% sucrose solution droplets directly to the mouthparts and left for 18 hours (overnight) in a separate controlled environment room under identical conditions as the rearing room. For unknown reasons, 24 workers did not survive overnight and were excluded from any data analysis. Between 08:00-09:00 the PER testing began on the remaining bees (n=413) by first testing their PER responsiveness to a 50% sucrose solution. Immediately after, each bee was fed a small droplet (0.8µl) of the sucrose solution for motivation 15 minutes before the start of the PER test (Figure S1). PER conditioning was conducted in front of a filtered ventilation system (Expo Drills & Tools AB500 Extractor fan), preventing the odour coming in to contact with neighbouring harnessed bees. Each bee was initially conditioned by exposure to clean air for 5 seconds, followed by scented air for 10 seconds. A harnessed bee was positioned 3 cm away from a glass odour tube, with the airflow delivered at a constant rate of 80 ml/second (Tetra APS – 100), which was channelled through either a ‘clean’ unscented odour tube or diverted through a ‘scented’ odour tube containing a piece of filter paper (5 x 20 mm) impregnated with 1 µl of lemon essential oil (Naturally Thinking Ltd.). Airflow between the clean and scented tube was controlled by a solenoid valve (Nass Magnet 108-030-0257 24vAC/12vDC) connected to a Raspberry Pi 2 (Model B) computer to ensure each bee was exposed to a consistent amount of clean and scented air. To develop an association between the lemon odour and the reward, we touched the bee’s antennae with a droplet of 0.8 µl of 50% sucrose 6 seconds into the 10 second odour delivery phase and allowed the bee to feed.

Following trial 1, the odour and reward presentation sequence was repeated to each adult an additional nine times. The inter-trial interval (ITI) per individual was 10 minutes allowing us to conduct the PER testing in batches of up to 20 workers. We waited 15 seconds after the odour and reward presentation sequence before moving to the next individual (Smith and Burden, 2014). We recorded whether the bees showed a PER to the odour stimulus prior to or after the reward, which were defined as a learnt or non-learnt response respectively. This provided a number of learnt responses achieved by each worker over the nine trials, enabling us to estimate the probability per trial of workers demonstrating a learnt response for each treatment. If a bee responded to the initial conditioning trial (trial 1) before the reward had been presented (n=24) the individual was excluded from the experimental analyses. If a bee showed no PER (did not feed) even after the reward was provided, and exhibited this over the next three consecutive trials, the individual was removed from testing from that point and categorised as a non-learner.

### Micro-CT scanning

Linking variation in learning to differences in brain growth requires high-resolution imaging technology that can explore minute changes to soft tissue. Using traditional histological methods would have been technically challenging as it relies on physical extraction from the headcase followed by tissue fixation, dehydration, embedding and sectioning (Simmons and Swanson, 2009) increasing the risk of destructive sampling. Potentially this could have caused a greater change to neuropil volume than the experimental treatment itself. We attempted to overcome such challenges by using micro-computed tomography (µ-CT) scanning.

Following the PER assay, bees were humanely sacrificed by swiftly decapitating the live individual using a disposable surgery scalpel and heads immediately fully submerged in a 70/30% ethanol/de-ionised water solution in separate 1.5ml centrifuge tubes and stored at 5°C. Preparation of the heads followed precisely the published protocol by Smith et al. (2016) with the soft brain tissue being stained for seven days with phosphotungstic acid (PTA) before being CT scanned at a voxel size of 3.5 - 4 µm using a Nikon Metrology HMX ST 225 system (Nikon Metrology, Tring, UK). The staining and scanning methodology we employed has been shown to give us confidence in the accuracy of our measurements of these complex neuropil structures (Smith et al., 2016). The raw µCT data for each brain scan was reconstructed using CTPro 2.1 software (Nikon Metrology, Tring, UK) and processed using VG Studio Max 2.1 (Volume Graphics GmbH, Heidelberg, Germany). Each 3D reconstructed scan was then re-oriented to the same optimum plane-of-view for visualization, and for the neuropils in question we re-sliced into a new series of 2D images. For each sample, scan images were exported as 8-bit BMP image series at a standardized voxel size of 4 µm. In total, 92 worker brains were µCT scanned, but based on staining quality and that both left and right structures could be segmented (including both medial and lateral calyx for the mushroom body calyces) we successfully segmented the mushroom bodies for 78 workers, central body for 88, antennal lobes for 89, medullas for 71 and lobulas for 71 (Table S4). For the purposes of comparing relative volumes (absolute volume divided by ITD to correct for body size), bees were originally sampled to have a balanced representation across treatments and age but blind of learning performance.

### Neuropil volume measurements

Segmentation and volume analysis of brain structures was carried out using the software SPIERS 2.20 (Serial Paleontological Image Editing and Rendering System). For segmentation, scan slices were converted to binary threshold images (of white active pixels and black inactive pixels) adjusted to achieve an optimum ratio of active white pixels that comprise the structure of interest, and inactive black pixels for the surrounding tissue. For each component structure, looped splines were placed around the active pixels at regular five slice intervals which were then interpolated across all slices between the intervals, so that each structure could be defined as an independent object for 3D reconstruction and volumetric calculation (for full segmentation protocol see Smith et al. 2016). This soft tissue segmentation protocol has been shown to provide repeatable and precise volumetric measurements of morphological structures of the bumblebee brain. To calculate absolute volume of each structure we used the voxel count function in SPIERS Edit, with relative volume calculated by dividing by the inter-tegula width (standard proxy for body size (Cane, 1987)). Inter-tegula width was measured using digital callipers (Workzone®) with the mean of two repeated measurements used. A single value was used in our analyses for each of the mushroom bodies, antennal lobes, lobulas and medullas by summing the volume of the left and right paired structures.

### Data Analysis

Statistical analyses were conducted in R version 3.5.1 (R Development Core Team 2018) using RStudio version 1.1.463, with models implemented using the lme4 package (Bates et al., 2015). For all models we included *treatment* as a fixed categorical factor. We considered measure of bee body size (*ITD*) as a continuous variable and *colony* as a random factor in our models if inclusion increased the fit of the model (model comparisons were assessed by comparing the AIC) otherwise they were not retained. For responsiveness and learning the data was analysed using the proportion of individuals showing a response with a generalized linear model (glm) using a binomial distribution and included the categorical variable *age* (3 or 12-day) as an additional fixed factor with *age* x *treatment* interaction term removed as it showed no significant effect. For looking at the proportion of learners by trial we used a linear mixed effects model (lmer) in which treatment consisted of two categories, *control* workers and pesticide workers (pooled from all three pesticide treatments due to low sample sizes per treatment). We considered a second order polynomial fit for *trial* number and individual *ID* as a random factor. For relative neuropil volumes we used a linear mixed-effects model (LMER) that included *age, ITD* and *colony* (random effect). We used a binomial generalised linear model (glm) to analyse how calyces’ volume influenced the learning score, as a proportion of the maximum learning score that could be achieved using calyces volume to analyse the *calyces volume* x *treatment* interaction on score.

## Author Contributions

RJG conceived the project; DBS & RJG designed the experiment; DBS, ARR & PHB conducted the experiment; DBS & FA carried out the µCT scanning; DBS & DB reconstructed and segmented the brains; ANA, DBS & RJG performed data analyses; DBS & RJG wrote the manuscript and ANA provided critical feedback.

## Acknowledgements

We thank Russell Garwood for help using SPIERS software, Dan Sykes and Amin Garbout for help with the µCT scanning protocol, Paul Beasley for technical support, Alfredo Sánchez Tójar and Peter Graystock for advice on da ta analysis, and Richard Abel, Mark Brown, Inti-Pedroso, Nigel Raine and Seirian Sumner for advice on pilot work. This work was supported by NERC grants (NE/L00755X/1 & NE/P012574/1) awarded to RJG which funded ANA and ARR. DBS’s PhD was supported by a NERC funded SSCP DTP scholarship in affiliation with the Grantham Institute at Imperial College London. RJG is supported by Imperial College’s Grand Challenges in Ecosystems and the Environment initiative.

## Supplementary Figures

**Figure S1.**
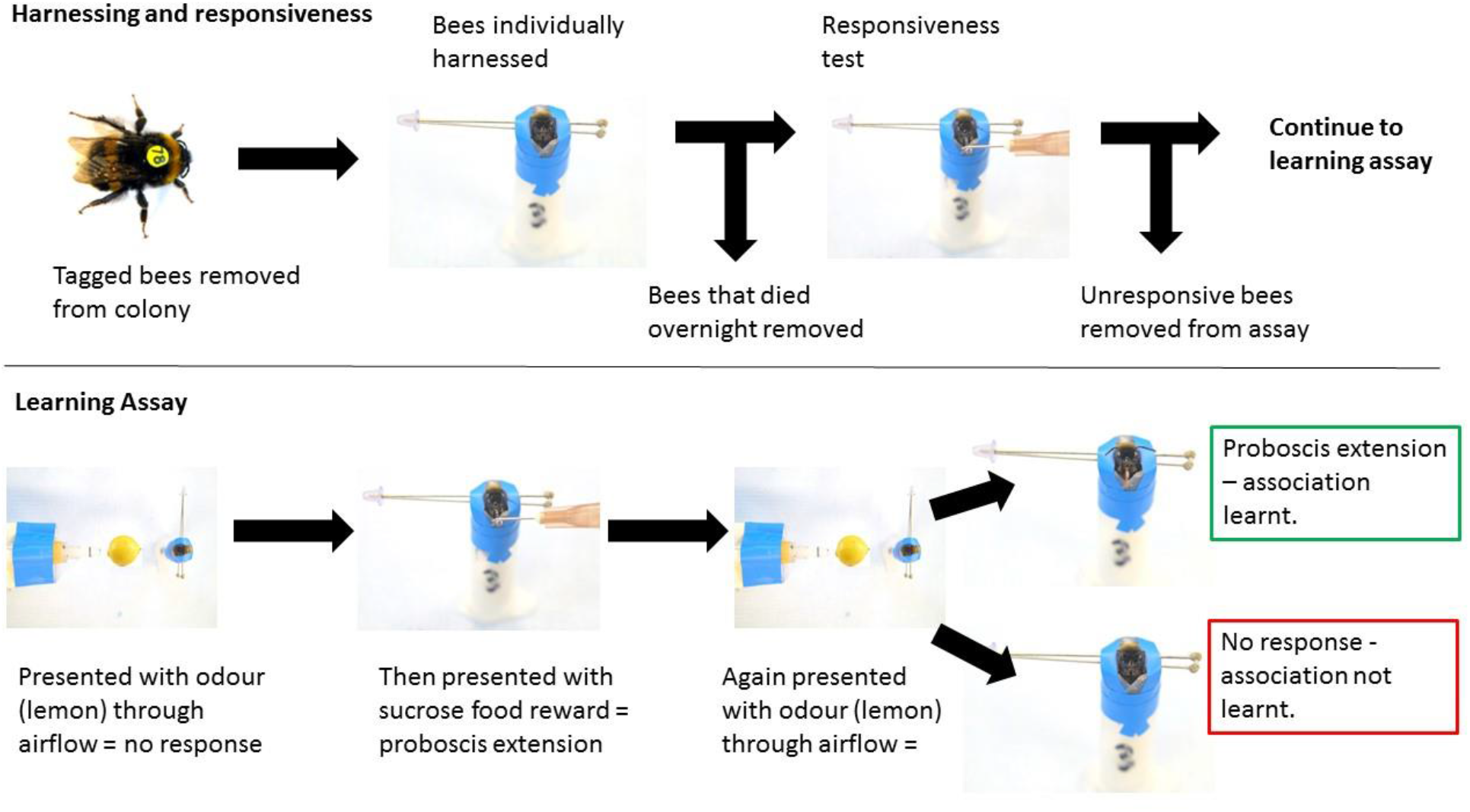
Proboscis extension reflex assay setup and step-by-step guide. Individuals were placed inside a plastic test tube and the tube was placed on ice for 10 minutes. Individuals were then harnessed in modified 2ml centrifuge tubes and a split pin yoke held them in place with electrical tape (blue). Harnessed bees were always placed the same distance from the air flow odour source and an extractor fan was mounted behind to remove excess odour.

**Figure S2.**
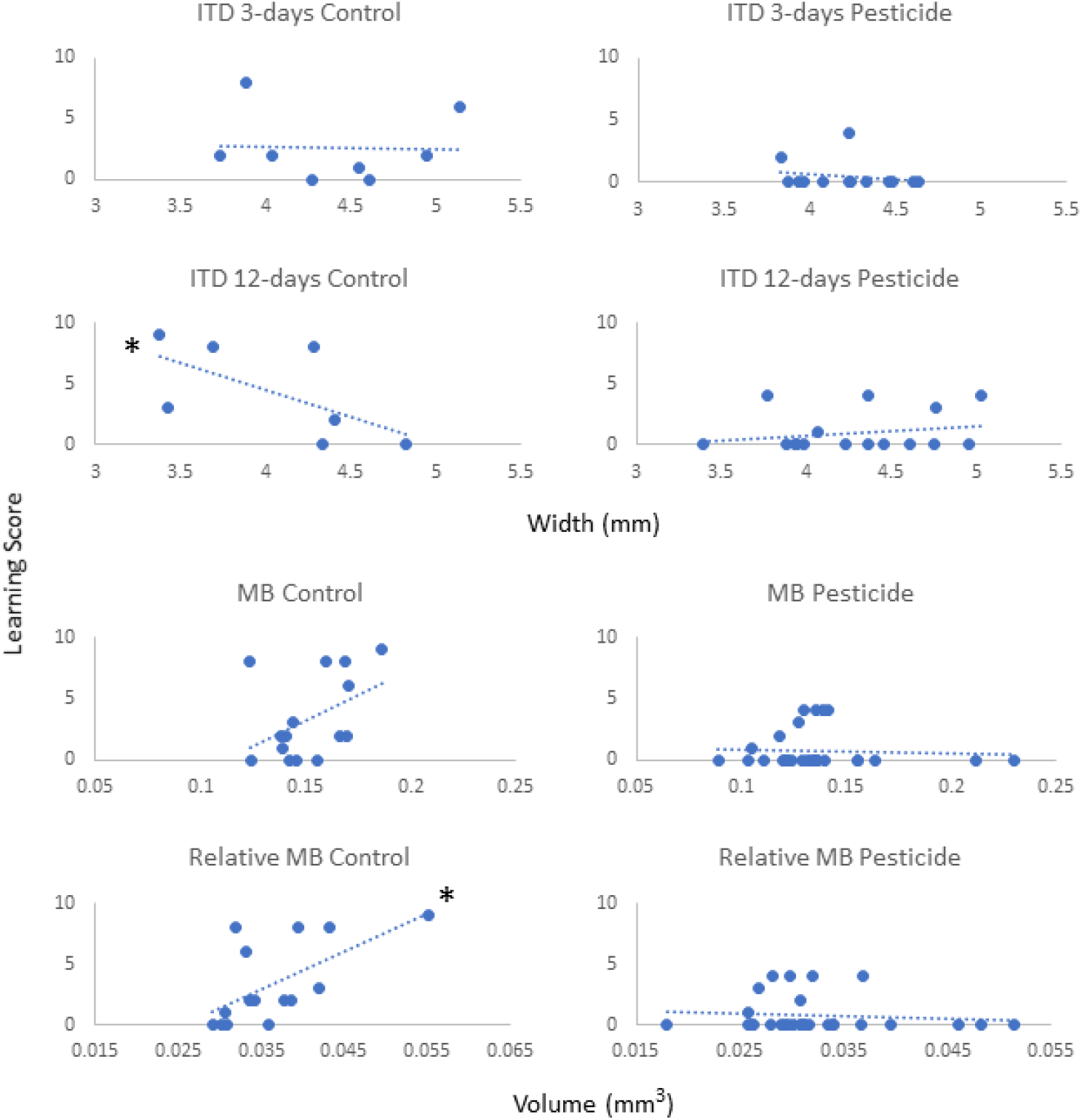
Predictor variables individual body size and mushroom body (MB) volumes were plotted against respective learning scores. The effect of width (mm) of the intertegula distance (ITD; proxy for body size) for 3-day and 12-day adult workers are shown. The absolute and relative (corrected for body size) whole mushroom body volumes (mm^3^), representing the combined volumes of the left and right hemispheres, for all workers regardless of age are shown. Filled circles represent the raw data with a dashed trend line fitted. Asterisks denote a significant negative or positive relationship (α=0.05) based on a Spearman’s Rank correlation analysis (detailed in Table S8).

**Figure S3.**
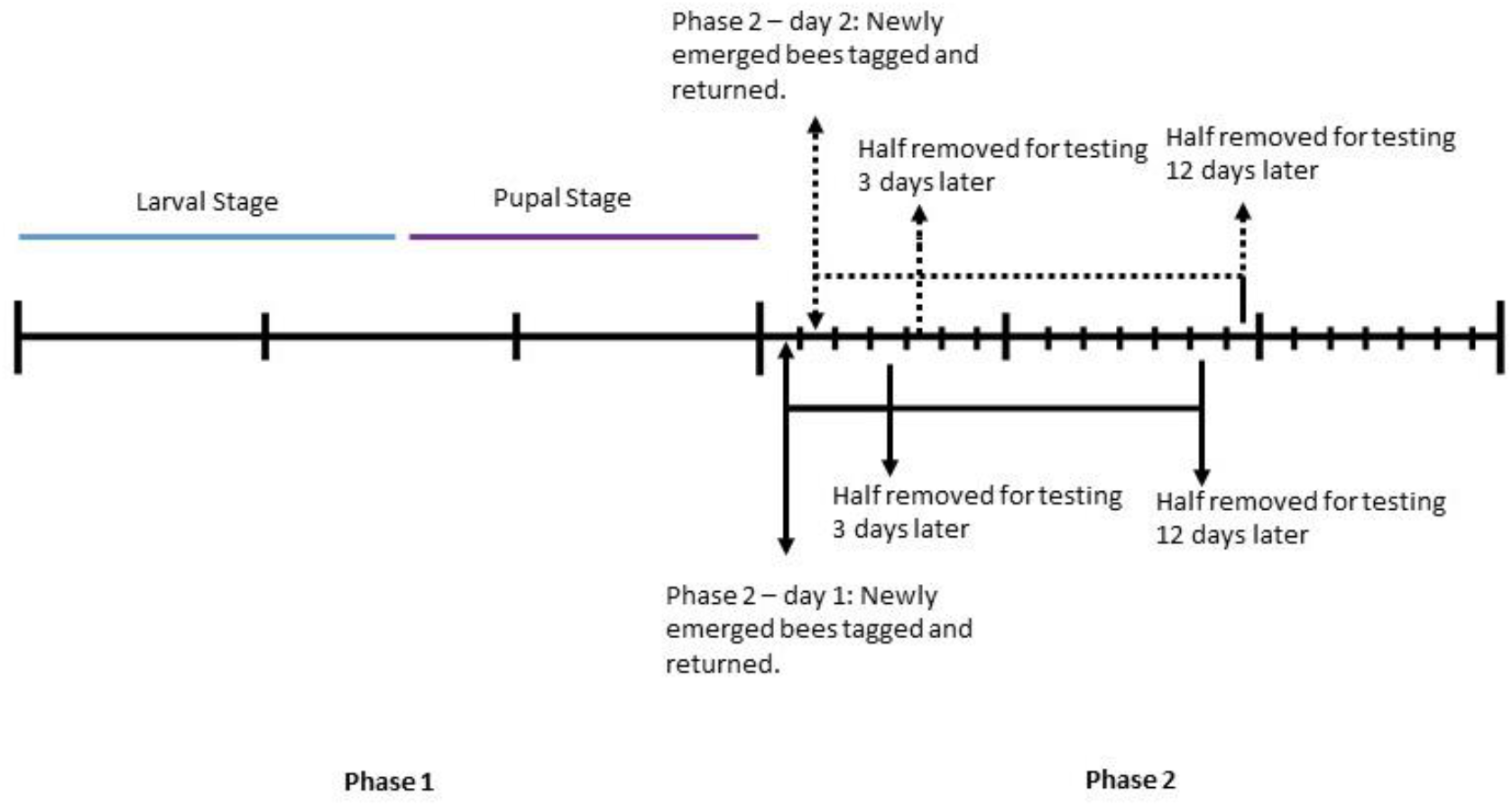
Sampling for 3 and 12-day adult cohorts. During Phase I: all newly eclosed bees were marked using a white paint pen, but this also represents the developmental time of workers with our sampled bees having been larvae (blue) and pupae (purple) during this 21 day development period. Phase II: for 11 days, colonies were checked daily with any newly eclosed workers being tagged and returned to their natal colony with half randomly assigned to a 3-day cohort and the other half a 12-days cohort. The respective cohorts were then removed 3 days or 12 days later. This continual tagging and sampling over the 11 day period provided us with a large number of workers to test (here we provide an example for the first two days for demonstration purposes).

## Supplementary Tables

**Table S1.**
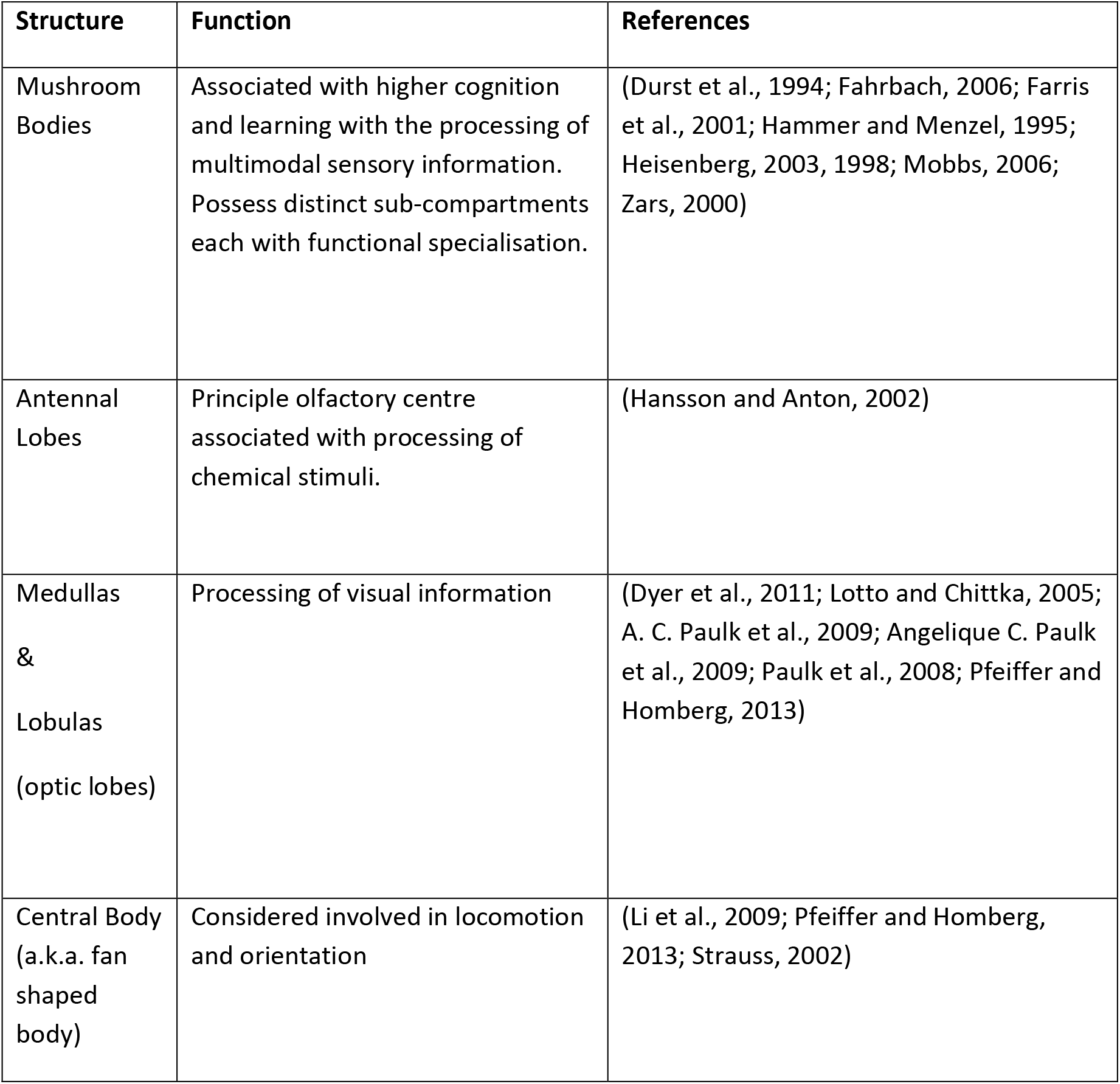
Informed from honeybee studies, our study focused on five key neuropils considered to be involved in the following primary functional roles.

| Structure | Function | References |
| --- | --- | --- |
| Mushroom Bodies | Associated with higher cognition and learning with the processing of multimodal sensory information. Possess distinct sub-compartments each with functional specialisation. | (Durst et al., 1994; Fahrbach, 2006; Farris et al., 2001; Hammer and Menzel, 1995; Heisenberg, 2003, 1998; Mobbs, 2006; Zars, 2000) |
| Antennal Lobes | Principle olfactory centre associated with processing of chemical stimuli. | (Hansson and Anton, 2002) |
| Medullas & Lobulas (optic lobes) | Processing of visual information | (Dyer et al., 2011; Lotto and Chittka, 2005; A. C. Paulk et al., 2009; Angelique C. Paulk et al., 2009; Paulk et al., 2008; Pfeiffer and Homberg, 2013) |
| Central Body (a.k.a. fan shaped body) | Considered involved in locomotion and orientation | (Li et al., 2009; Pfeiffer and Homberg, 2013; Strauss, 2002) |

**Table S2.**
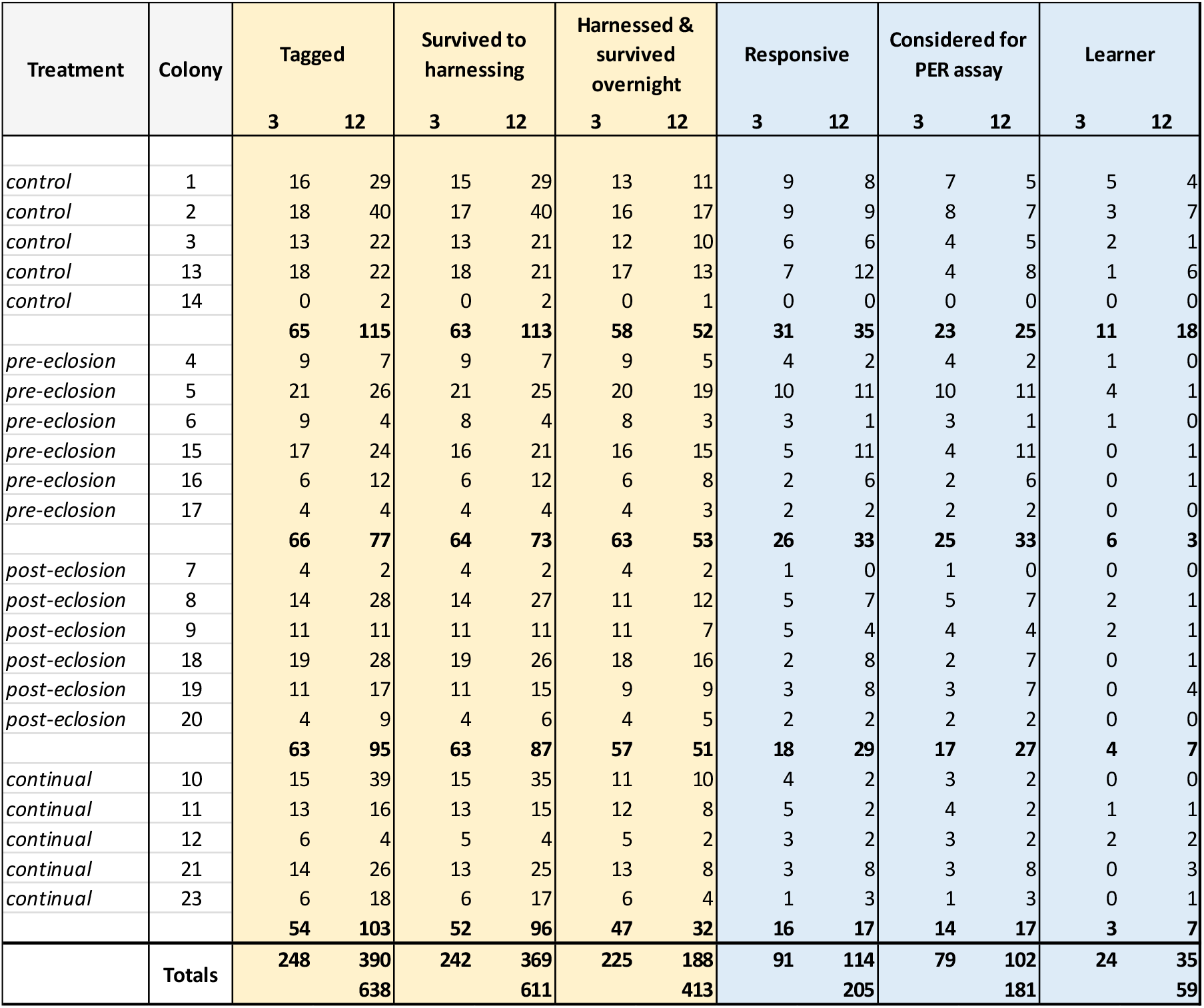
Number of 3- and 12-day adult workers from each colony that were prepared for tested using our PER setup. ‘Tagged’ = number of newly emerged workers that were tagged with a unique colour and numbered Opalith tag; ‘Survived to Harnessing’ = number of tagged workers survived to 3- or 12-days post-emergence and could be harnessed for the PER assay; ‘Harnessed & survived overnight’ = number of workers that were harnessed but also were alive inside the harness the next day (n=24 died overnight); ‘Responsive’ = number of workers that exhibited a PER response on touching the antenna with a 50% sucrose solution droplet; ‘Considered for PER assay’ = the number of workers remaining once any workers showing a PER to the lemon odour and before the sucrose provision on the 1^st^ trial were removed; ‘Learner’ = number of workers that showed at least one olfactory conditioned PER response over the ten trials.

**Table S3.** Statistical comparisons of responsiveness and learning. **a-b)** statistical outputs from binomial Generalized Linear Model in R (GLM) when analysing the proportion of workers that showed a PER response to a sucrose droplet prior to undertaking the learning assay, and the proportion of workers showing at least one olfactory conditioned PER learnt response during trials 2-10 of the learning assay. Exposure treatments are comparisons to *control* workers (‘intercept’) with *age* (3 versus 12-day workers) and ITD (inter-tegula distance which is a proxy for body size) considered. For both models the interaction term between pesticide treatment and age was removed as no significant effect could be detected. **c)** statistical output from a binomial Linear Mixed Effects Model in R (LMER) when analysing the proportion of workers that showed a PER response over the PER assay trials. For this model all workers from the three pesticide exposure treatments were pooled to compare one cohort against *control* workers (intercept). A 2^nd^ order polynomial relationship was found to be the best fit to the data. Significant differences (α=0.05) are highlighted in bold.

| **a) Responsive** |  |  |  |  |
| --- | --- | --- | --- | --- |
| GLM(responsive(y/n) ~ treatment + age + ITD, family = binomial) |  |  |  |  |
|  | Estimate | Std. Error | z value | Pr(>\|z\|) |
| (Intercept) | -1.279 | 0.959 | -1.333 | 0.182 |
| pre-eclosion | -0.342 | 0.276 | -1.242 | 0.214 |
| post-eclosion | -0.714 | 0.282 | -2.529 | **0.011** |
| continual | -0.737 | 0.307 | -2.400 | **0.016** |
| age | 0.842 | 0.205 | 4.101 | **<0.001** |
| ITD | 0.306 | 0.218 | 1.406 | 0.160 |
| **b) Learners** |  |  |  |  |
| GLM(learner(y/n) ~ treatment + age + ITD, family = binomial) |  |  |  |  |
| (Intercept) | -3.865 | 1.820 | -2.123 | 0.034 |
| pre-eclosion | -2.086 | 0.476 | -4.382 | **<0.001** |
| post-eclosion | -1.663 | 0.476 | -3.490 | **<0.001** |
| continual | -1.409 | 0.507 | -2.778 | **0.005** |
| age | 0.413 | 0.360 | 1.147 | 0.251 |
| ITD | 0.953 | 0.413 | 2.308 | **0.021** |
| **c) Learners by trial** |  |  |  |  |
| LMER(learner(y/n) ~ treatment + poly(trial,2) + (1\|worker_id), family = binomial) |  |  |  |  |
| (Intercept) | -0.889 | 0.308 | -2.887 | 0.004 |
| pesticide_treatments | -0.861 | 0.413 | -2.083 | **0.037** |
| poly(trial, 2)1 | 48.427 | 5.398 | 8.972 | **<0.001** |
| poly(trial, 2)2 | -18.909 | 3.965 | -4.769 | **<0.001** |

**Table S4.**
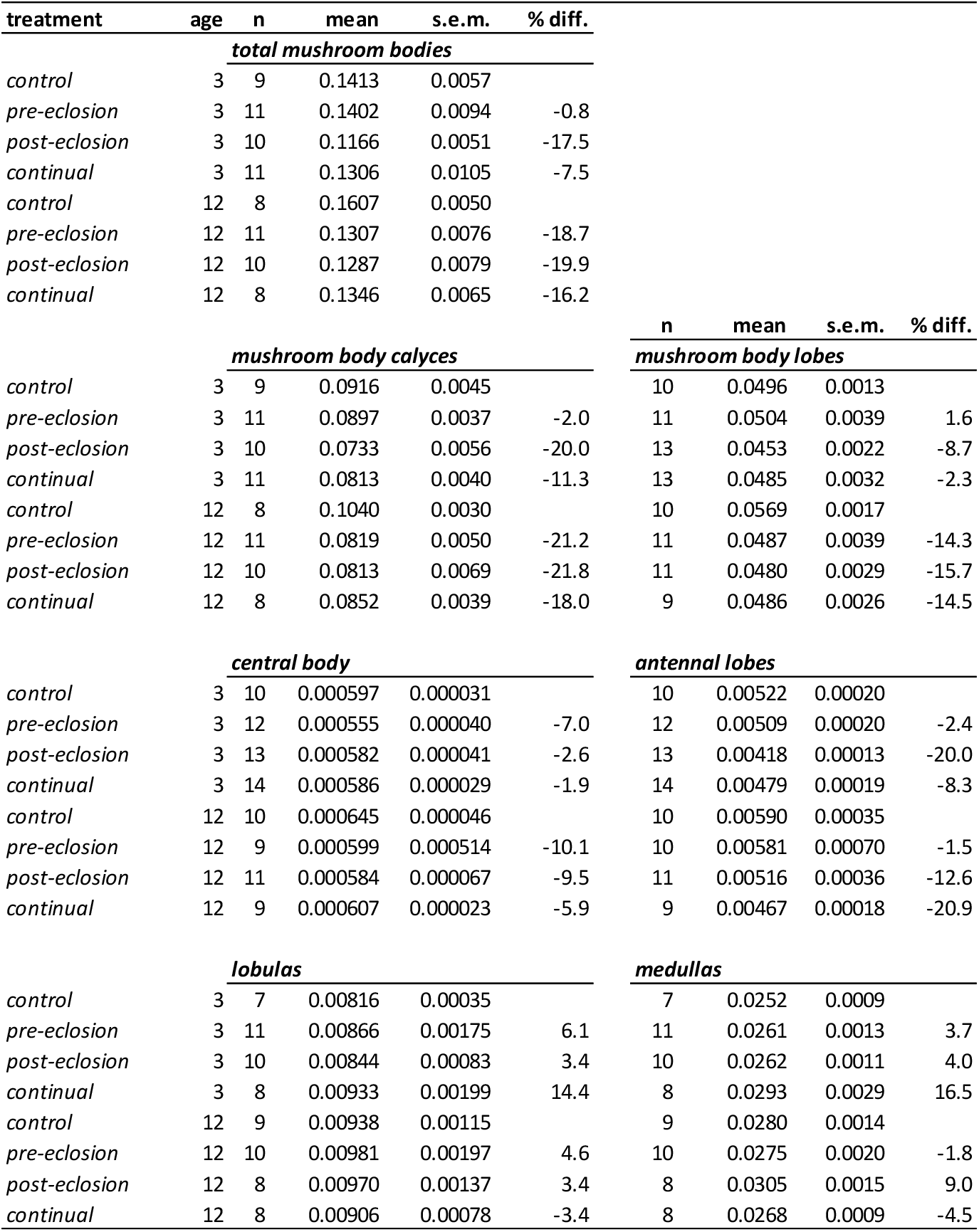
Average volumes (mm^3^) of segmented brain neuropils using µCT scanning for each experimental treatment and age cohort. Mean (± s.e.m.) values are based on workers across all colonies and represent relative volumes (absolute volume divided by workers size). The percentage difference (% diff.) of the mean of each pesticide treatment relative to the control group is provided with negative values showing smaller and positive values showing larger average volumes. Based on the quality of staining and scanning, sample sizes (n) for each neuropil were limited to those that had both structures from the left and right brain hemispheres successfully segmented. (N.B. for the *central body* one individual (18G) was removed from the analysis as it represented an extreme outlier likely caused by a segmentation error).

**Table S5.** Statistical comparisons of mushroom body relative volumes. Statistical outputs from a Generalized Mixed Effects Model in R (LMER) for the combined volumes of the left and right **a.** whole mushroom bodies, **b.** the medial and lateral calyces of the mushroom body together, **c.** lobes of the mushroom body for only those individuals that also had the calyxes segmented (n=78), d. the lobes of the mushroom body for any individual that had them segmented (n=88). Exposure treatments are comparisons to *control* workers (‘intercept’) with *age* (3 versus 12-day workers) and ITD (inter-tegula distance - proxy for body size) considered. For both models the interaction term between pesticide treatment and age was removed as no significant effect could be detected. Significant differences (α=0.05) highlighted in bold black.

| **a) Total Mushroom Bodies (n=78)** |  |  |  |  |  |
| --- | --- | --- | --- | --- | --- |
| LMER(relativeMB ~ treatment + age + size + (1\|colony)) |  |  |  |  |  |
|  | Estimate | Std. Error | df | t value | Pr(>\|t\|) |
| (Intercept) | 0.058 | 0.008 | 43.244 | 7.732 | <0.001 |
| *pre-eclosion* | -0.004 | 0.002 | 5.845 | -1.822 | 0.120 |
| *post-eclosion* | -0.007 | 0.002 | 6.166 | -2.873 | **0.027** |
| *continual* | -0.005 | 0.002 | 6.348 | -2.032 | 0.086 |
| *age* | 0.002 | 0.001 | 67.913 | 1.473 | 0.145 |
| *ITD* | -0.006 | 0.002 | 49.038 | -3.246 | **0.002** |
| **b) Total Mushroom Body Calyces (n=78)** |  |  |  |  |  |
| LMER(relativeMBCalyces ~ treatment + age + size + (1\|colony)) |  |  |  |  |  |
| (Intercept) | 0.035 | 0.005 | 40.979 | 7.735 | <0.001 |
| *pre-eclosion* | -0.003 | 0.001 | 6.637 | -2.409 | **0.049** |
| *post-eclosion* | -0.005 | 0.001 | 7.212 | -3.828 | **0.006** |
| *continual* | -0.004 | 0.001 | 7.626 | -2.896 | **0.021** |
| *age* | 0.001 | 0.001 | 69.152 | 1.565 | 0.122 |
| *ITD* | -0.003 | 0.001 | 44.961 | -2.919 | **0.005** |
| **c) Total Mushroom Body Lobes (n=78)** |  |  |  |  |  |
| LMER(relativeMBLobes ~ treatment + age + size + (1\|colony)) |  |  |  |  |  |
| (Intercept) | 0.024 | 0.003 | 65.458 | 8.182 | <0.001 |
| *pre-eclosion* | -0.001 | 0.001 | 6.440 | -0.890 | 0.406 |
| *post-eclosion* | -0.001 | 0.001 | 6.184 | -1.181 | 0.281 |
| *continual* | -0.001 | 0.001 | 6.350 | -0.936 | 0.383 |
| *age* | 0.001 | 0.000 | 74.263 | 1.137 | 0.259 |
| *ITD* | -0.003 | 0.001 | 76.887 | -4.285 | **<0.001** |
| **d) Total Mushroom Body Lobes (n=88)** |  |  |  |  |  |
| LMER(relativeMBLobes ~ treatment + age + size + (1\|colony)) |  |  |  |  |  |
| (Intercept) | 0.024 | 0.003 | 53.979 | 7.295 | <0.001 |
| *pre-eclosion* | -0.001 | 0.001 | 5.995 | -0.865 | 0.421 |
| *post-eclosion* | -0.002 | 0.001 | 6.217 | -1.387 | 0.213 |
| *continual* | -0.001 | 0.001 | 6.261 | -0.757 | 0.476 |
| *age* | 0.001 | 0.001 | 66.485 | 1.145 | 0.256 |
| *ITD* | -0.003 | 0.001 | 63.112 | -3.661 | **<0.001** |

**Table S6.** Statistical comparisons of the relative volumes of the central body, antennal lobes, lobulas and medullas. Statistical outputs are for the combined volumes of the left and right (except the central body), from a Generalized Mixed Effects Model in R (LMER). Exposure treatments are comparisons to *control* workers (‘intercept’) with *age* (3 versus 12-day workers) and ITD (inter-tegula distance which is a proxy for body size) considered. For both models the interaction term between pesticide treatment and age was removed as no significant effect could be detected. Significant differences (α=0.05) highlighted in bold black.

| Central Body n=88 |  |  |  |  |  |
| --- | --- | --- | --- | --- | --- |
| lmer - central Body ~ treatment + age + size + (1\|colony) |  |  |  |  |  |
|  | Estimate | s.e. | df | t | p |
| (Intercept) | 0.000573 | 0.000176 | 64.7 | 3.26 | 0.002 |
| Pre-eclosion | 0.000074 | 0.000073 | 5.9 | 1.01 | 0.351 |
| Post-eclosion | -0.000002 | 0.000072 | 5.6 | -0.03 | 0.976 |
| Continual | 0.000001 | 0.000072 | 5.7 | 0.02 | 0.985 |
| age | 0.000027 | 0.000028 | 74.4 | 0.96 | 0.339 |
| size | -0.000102 | 0.000038 | 78.4 | -2.67 | **0.009** |
| Antennal Lobes n=89 |  |  |  |  |  |
| lmer - antennal lobes ~ treatment + age + size + (1\|colony) |  |  |  |  |  |
| (Intercept) | 0.003658 | 0.000320 | 63.6 | 11.42 | <0.001 |
| Pre-eclosion | -0.000034 | 0.000133 | 5.5 | -0.26 | 0.807 |
| Post-eclosion | -0.000208 | 0.000130 | 5.3 | -1.60 | 0.168 |
| Continual | -0.000176 | 0.000131 | 5.3 | -1.35 | 0.232 |
| age | 0.000101 | 0.000051 | 73.9 | 1.98 | 0.051 |
| size | -0.000553 | 0.000070 | 77.9 | -7.96 | **<0.001** |
| Lobulas n=71 |  |  |  |  |  |
| lmer - lobulas ~ treatment + age + size + (1\|colony) |  |  |  |  |  |
| (Intercept) | 0.004984 | 0.000493 | 47.1 | 10.11 | <0.001 |
| Pre-eclosion | 0.000052 | 0.000140 | 6.4 | 0.37 | 0.723 |
| Post-eclosion | 0.000034 | 0.000142 | 7.3 | 0.24 | 0.818 |
| Continual | 0.000057 | 0.000146 | 7.4 | 0.39 | 0.706 |
| age | 0.000192 | 0.000083 | 61.5 | 2.30 | **0.025** |
| size | -0.000693 | 0.000108 | 52.2 | -6.44 | **<0.001** |
| Medullas n=71 |  |  |  |  |  |
| lmer - medullas ~ treatment + age + size + (1\|colony) |  |  |  |  |  |
| (Intercept) | 0.014020 | 0.001639 | 42.7 | 8.55 | <0.001 |
| Pre-eclosion | -0.000094 | 0.000425 | 6.7 | -0.22 | 0.832 |
| Post-eclosion | 0.000295 | 0.000435 | 8.2 | 0.68 | 0.517 |
| Continual | 0.000208 | 0.000447 | 8.0 | 0.46 | 0.655 |
| age | 0.000369 | 0.000287 | 62.5 | 1.29 | 0.203 |
| size | -0.001826 | 0.000360 | 46.9 | -5.08 | **<0.001** |

**Table S7.** Statistical output from a binomial generalised linear model in R (GLM) when analysing the final learning score achieved by each worker by the end of PER assay trials. For this model all workers from the three pesticide exposure treatments were pooled to compare one cohort against *control* workers (intercept). Significant differences (α=0.05) are highlighted in bold.

| glm(learning_score ~ relativeMBCalyces*treatment, family = binomial) |  |  |  |  |
| --- | --- | --- | --- | --- |
|  | Estimate | Std. Error | z value | Pr(>\|z\|) |
| (Intercept) | -6.992 | 1.416 | -4.939 | <0.001 |
| calyces | 265.823 | 58.981 | 4.507 | **<0.001** |
| pesticide | 5.611 | 1.802 | 3.113 | **0.002** |
| calyces*pesticide | -321.899 | 81.268 | -3.961 | **<0.001** |

**Table S8.**
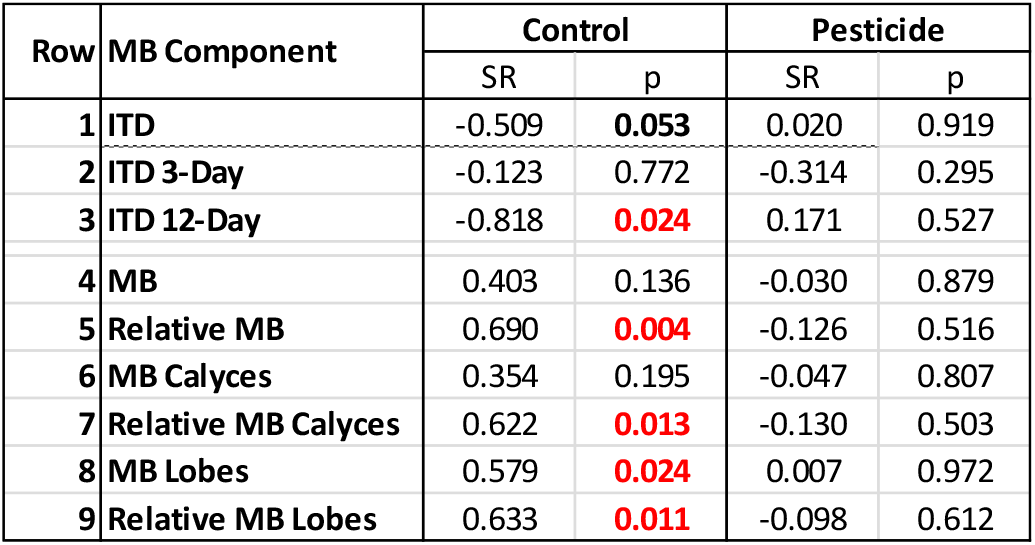
Spearman’s rank correlations to investigate the relationship between the response variable learning score and the predictor variables body size (ITD) or mushroom body volume. Row 1 is intertegula distance (ITD; proxy for body size) for workers regardless of age; Rows 2 & 3 are ITDs for 3 and 12-day workers (respectively); Rows 4-5 is the absolute volume of the whole mushroom bodies (left and right combined), and relative volume of the whole mushroom bodies (left and right combined) corrected for body size, respectively; Rows 6-9 also show absolute and relative volumes for the calyces and lobes separately (left and right combined). Significant differences (α=0.05) are highlighted in bold red, with near significant differences (α=0.1) in bold black.

| Row | MB Component | Control |  | Pesticide |  |
| --- | --- | --- | --- | --- | --- |
|  |  | SR | p | SR | p |
| 1 | ITD | -0.509 | **0.053** | 0.020 | 0.919 |
| 2 | ITD 3-Day | -0.123 | 0.772 | -0.314 | 0.295 |
| 3 | ITD 12-Day | -0.818 | **0.024** | 0.171 | 0.527 |
| 4 | MB | 0.403 | 0.136 | -0.030 | 0.879 |
| 5 | Relative MB | 0.690 | **0.004** | -0.126 | 0.516 |
| 6 | MB Calyces | 0.354 | 0.195 | -0.047 | 0.807 |
| 7 | Relative MB Calyces | 0.622 | **0.013** | -0.130 | 0.503 |
| 8 | MB Lobes | 0.579 | **0.024** | 0.007 | 0.972 |
| 9 | Relative MB Lobes | 0.633 | **0.011** | -0.098 | 0.612 |

**Table S9.**
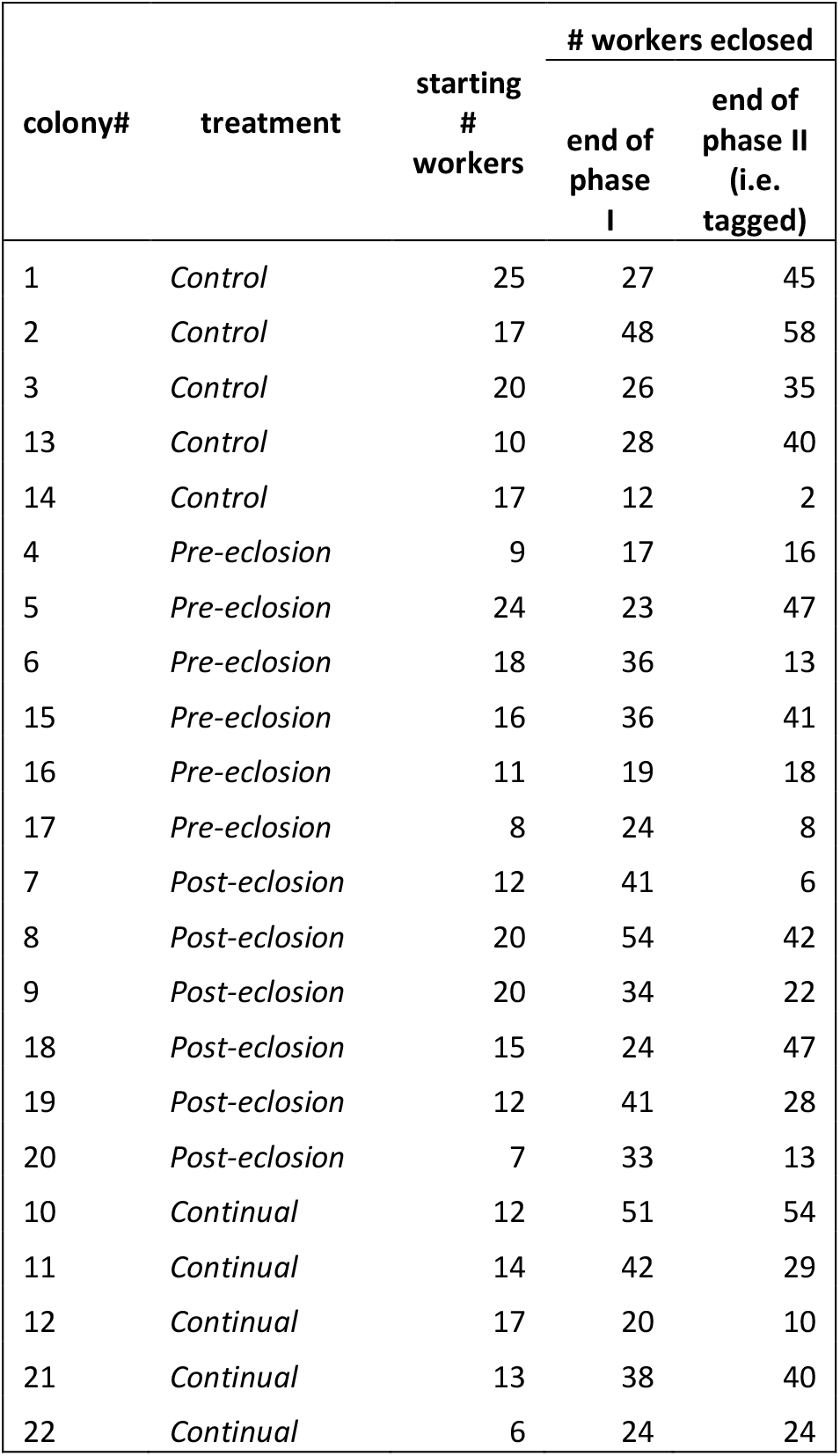
Colony census at start of the experiment, and at the end of each phase. All workers eclosing during Phase II were tagged with a unique colour and numbered Opalith tag so the age of each worker was known.

**Table S10.** Colony daily sucrose consumption (ml) as the experiment progressed (days 1-43). Of the total provisioned sucrose (1,110ml per colony), *control, pre-eclosion, post-eclosion* and *continual* treatments consumed a median (IQR) of 54 (50-56), 47 (37-57), 61 (48-69) and 51 (35-63) % respectively.

| Treatment | Colony # | Consumption (ml) per day |  |  |  |  |  |  |  |  |  |  |  |  |  |  |  |  |  |  |  |  |  |
| --- | --- | --- | --- | --- | --- | --- | --- | --- | --- | --- | --- | --- | --- | --- | --- | --- | --- | --- | --- | --- | --- | --- | --- |
|  |  | 1 | 3 | 5 | 7 | 9 | 11 | 13 | 15 | 17 | 19 | 21 | 23 | 25 | 27 | 29 | 31 | 33 | 35 | 37 | 39 | 41 | 43 |
| Control | 1 | 9 | 15 | 25 | 15 | 18 | 21 | 20 | 22 | 28 | 34 | 15 | 40 | 30 | 35 | 37 | 37 | 31 | 36 | 35 | 33 | 33 | 37 |
| Control | 2 | 14 | 20 | 29 | 17 | 20 | 20 | 29 | 28 | 27 | 33 | 13 | 28 | 28 | 30 | 30 | 34 | 26 | 32 | 35 | 38 | 37 | 35 |
| Control | 3 | 10 | 10 | 19 | 14 | 20 | 20 | 21 | 22 | 23 | 30 | 14 | 30 | 28 | 38 | 37 | 38 | 30 | 37 | 35 | 37 | 39 | 39 |
| Pre-eclosion | 4 | 7 | 10 | 13 | 10 | 13 | 14 | 15 | 12 | 16 | 19 | 9 | 20 | 17 | 20 | 16 | 17 | 16 | 21 | 21 | 22 | 25 | 26 |
| Pre-eclosion | 5 | 13 | 12 | 24 | 18 | 24 | 22 | 20 | 20 | 23 | 26 | 12 | 27 | 25 | 30 | 31 | 32 | 28 | 32 | 30 | 33 | 35 | 35 |
| Pre-eclosion | 6 | 25 | 21 | 31 | 24 | 33 | 33 | 31 | 31 | 29 | 39 | 14 | 41 | 34 | 39 | 37 | 40 | 28 | 35 | 37 | 35 | 37 | 45 |
| Post-eclosion | 7 | 13 | 10 | 29 | 13 | 17 | 18 | 16 | 17 | 20 | 26 | 12 | 26 | 37 | 45 | 46 | 43 | 36 | 43 | 48 | 48 | 54 | 59 |
| Post-eclosion | 8 | 29 | 22 | 32 | 21 | 24 | 24 | 29 | 23 | 30 | 37 | 14 | 30 | 34 | 39 | 39 | 38 | 32 | 42 | 57 | 42 | 45 | 52 |
| Post-eclosion | 9 | 18 | 17 | 30 | 23 | 28 | 28 | 35 | 39 | 35 | 39 | 21 | 46 | 42 | 48 | 53 | 52 | 36 | 46 | 52 | 48 | 55 | 55 |
| continual | 10 | 19 | 18 | 25 | 20 | 22 | 25 | 27 | 28 | 33 | 35 | 13 | 35 | 35 | 40 | 42 | 46 | 27 | 40 | 36 | 40 | 40 | 35 |
| continual | 11 | 20 | 15 | 22 | 24 | 25 | 25 | 28 | 26 | 28 | 32 | 15 | 35 | 35 | 39 | 36 | 46 | 36 | 38 | 35 | 35 | 33 | 32 |
| continual | 12 | 5 | 15 | 17 | 12 | 15 | 15 | 21 | 16 | 18 | 25 | 11 | 24 | 1 | 29 | 25 | 28 | 25 | 31 | 33 | 36 | 40 | 41 |
| Control | 13 | 17 | 17 | 17 | 23 | 21 | 25 | 21 | 26 | 24 | 28 | 15 | 32 | 30 | 36 | 33 | 35 | 38 | 40 | 39 | 40 | 40 | 41 |
| Control | 14 | 15 | 16 | 14 | 17 | 13 | 15 | 16 | 20 | 16 | 21 | 12 | 25 | 23 | 34 | 30 | 31 | 29 | 33 | 33 | 35 | 32 | 34 |
| Pre-eclosion | 15 | 17 | 19 | 23 | 26 | 25 | 29 | 24 | 29 | 25 | 29 | 17 | 37 | 28 | 33 | 31 | 31 | 30 | 30 | 26 | 31 | 29 | 31 |
| Pre-eclosion | 16 | 17 | 18 | 14 | 23 | 17 | 22 | 18 | 22 | 20 | 24 | 14 | 30 | 27 | 29 | 25 | 25 | 23 | 23 | 22 | 24 | 23 | 23 |
| Pre-eclosion | 17 | 13 | 14 | 11 | 19 | 16 | 21 | 19 | 22 | 18 | 20 | 11 | 25 | 21 | 23 | 23 | 23 | 23 | 23 | 21 | 21 | 17 | 19 |
| Post-eclosion | 18 | 15 | 15 | 15 | 22 | 18 | 22 | 29 | 30 | 19 | 24 | 14 | 28 | 23 | 27 | 36 | 32 | 30 | 28 | 32 | 25 | 27 | 27 |
| Post-eclosion | 19 | 23 | 23 | 23 | 29 | 25 | 31 | 26 | 36 | 34 | 36 | 21 | 38 | 33 | 38 | 29 | 34 | 34 | 36 | 36 | 34 | 34 | 33 |
| Post-eclosion | 20 | 15 | 15 | 20 | 26 | 20 | 23 | 22 | 26 | 21 | 26 | 13 | 23 | 22 | 26 | 25 | 29 | 30 | 31 | 28 | 27 | 26 | 26 |
| Continual | 21 | 18 | 19 | 19 | 26 | 25 | 29 | 29 | 37 | 30 | 39 | 21 | 42 | 35 | 39 | 41 | 43 | 45 | 46 | 46 | 47 | 45 | 46 |
| Continual | 22 | 13 | 13 | 10 | 13 | 10 | 12 | 9 | 10 | 7 | 10 | 4 | 10 | 7 | 12 | 12 | 12 | 12 | 13 | 15 | 16 | 9 | 11 |
| Continual | 23 | 14 | 15 | 15 | 21 | 13 | 16 | 16 | 19 | 16 | 19 | 12 | 24 | 22 | 25 | 25 | 25 | 23 | 25 | 24 | 25 | 24 | 26 |

